# Identification of human skin microbiome odorants that manipulate mosquito landing behavior

**DOI:** 10.1101/2023.08.19.553996

**Authors:** Iliano V. Coutinho-Abreu, Omid Jamshidi, Robyn Raban, Katayoon Atabakhsh, Joseph A. Merriman, Michael A. Fischbach, Omar S. Akbari

**Author notes:** To whom correspondence should be addressed: Omar S. Akbari, School of Biological Sciences, Department of Cell and Developmental Biology, University of California, San Diego, La Jolla, CA 92093, USA.

## Abstract

The resident human skin microbiome is responsible for the production of most of the human scents that are attractive to mosquitoes. Hence, engineering the human skin microbiome to synthesize less of mosquito attractants or produce repellents could potentially reduce bites and prevent the transmission of deadly mosquito-borne pathogens. In order to further characterize the human skin volatilome, we quantified the major volatiles of 39 strains of skin commensals (*Staphylococci* and *Corynebacterium*). Importantly, to validate the behavioral activity of these volatiles, we first assessed landing behavior triggered by human skin bacteria volatiles. We demonstrated that this behavioral step is gated by the presence of carbon dioxide and L-(+)-lactic acid, similar to the combinatorial coding triggering short range attraction. Repellency behavior to selected skin volatiles and the geraniol terpene was tested in the presence of carbon dioxide and L-(+)-lactic acid. In a 2-choice landing behavior context, the skin volatiles 2- and 3-methyl butyric acids reduced mosquito landing by 62.0-81.6% and 87.1-99.6%, respectively. Similarly, geraniol was capable of reducing mosquito landing behavior by 74.9%. We also tested the potential repellency effects of geraniol on mosquitoes at short-range using a 4-port olfactometer. In these assays, geraniol reduced mosquito attraction (69-78%) to a mixture of key human kairomones carbon dioxide, L-(+)-lactic acid, and ammonia. These findings demonstrate that carbon dioxide and L-(+)-lactic acid changes the valence of other skin volatiles towards mosquito landing behavior. Moreover, this study offers candidate odorants to be targeted in a novel strategy to reduce attractants or produce repellents by the human skin microbiota that may curtail mosquito bites, and subsequent mosquito-borne disease.

## Introduction

Mosquitoes are one of the biggest threats to human morbidity and mortality around the world due to their exceptional ability to transmit pathogens, including viruses, malaria parasites, and filarial worms. As the number of mosquito vectors resistant to commercial insecticides ^1^ and vector-borne pathogens gaining resistance to best-in-class drugs ^2,3^ has increased in recent years ^1^, innovative strategies to prevent mosquito bites and pathogen transmission are critical. Ideally, such strategies should protect against the bites of multiple mosquito vectors. Amongst potential new strategies to prevent mosquito bites are the development of safer, affordable, and globally accessible mosquito repellents ^4,5^. Current strategies aim to disrupt the mosquito chemosensory system using gene editing tools ^6–8^ and spreading these loss of function mutants into wild populations ^9^ with some success. As our understanding of attractive/repellent odorants and sources increases, alternative strategies may provide additional protection. With the human skin being the source of numerous attractive odorants, we investigate the impacts of reducing the production of attractive odorants and/or increasing the production of repellents by the human skin might also potentially reduce mosquito bites and pathogen transmission ^10,11^.

Synthetic mosquito repellents such as DEET and picaridin are effective at preventing mosquito bites ^12^. However, DEET can cause health issues ^12^, is unaffordable for widespread use ^13^, and requires reapplication within hours ^12^. In order to find alternative mosquito repellents, chemoinformatics ^4^ and machine learning approaches ^5^ have been used to interrogate chemical databases for molecules structurally similar to known repellents. A few of these candidate repellents have been shown to repel fruit flies ^14^; however, these candidates are yet to be shown effective against mosquitoes.

With the advent of genome editing, multiple mosquito chemosensory receptor genes have been modified to encode non-functional receptors, aiming at disrupting host seeking behavior. Genes encoding olfactory coreceptors orco ^6,15^, Ir25a ^16^, Ir76b ^16^, and Ir8a ^8^, carbon dioxide coreceptor Gr3 ^7,8,17^, heat receptors TripA1 ^18^, and Ir21a ^19^, and a humidity sensor co-receptor Ir93a ^20^ have been disrupted; nonetheless, these gene mutations were not sufficient to completely abrogate mosquito host seeking activity ^6–8,19^. Whether manipulating the activity of higher order neurons can effectively disrupt mosquito host seeking behavior ^21^, has yet to be determined.

The human scent emitted by the skin is produced by the microbiome resident in hair follicles and sweat glands^22^. Human sweat glands belong to three distinct classes, eccrine, apocrine, and sebaceous, which secrete amino acids, fatty acids, and salts, that are used as nutrients by the skin microbiome ^22^. The metabolization of these nutrients leads to the release of small molecules, such as L-(+)-lactic acid, ammonia, and short- and middle chain carboxylic acids ^23^, which synergizes with carbon dioxide in breath as well as body heat and humidity as attractants to anthropophilic mosquitoes ^10^. On the other hand, very little is known about how skin bacteria-derived volatiles drive mosquito landing behavior.

In order to unveil the hierarchical representation of skin volatiles that guide mosquito landing, and to identify natural odorants that prevent this fundamental step for mosquito blood feeding, we aimed to first address the contributions of skin commensals to mosquito behavior by 1) quantifying key metabolites/volatiles produced during growth in skin like media conditions, 2) determining the impact of a subset of skin commensal derived volatiles in *Aedes aegypti* landing behavior, and 3) evaluate a member of a known class of repellents to reduce *A. aegypti* attraction. These findings set the stage for the development of novel strategies to prevent mosquito bites through the manipulation of their olfactory system.

## Results

### Quantifying key volatiles produced by *Staphylococci* and *Corynebacterium* skin commensal isolates

Mosquito attractive volatiles originate from the human skin microbiome ^24^. *Staphylococci* and *Corynebacterium* are frequently amongst the top ten isolated and identified human skin commensals, with *Staphylococcus epidermidis* being considered one of 31 “core” human skin commensals around the world ^25^. By GC/MS, we sought to build a profile of these volatiles produced by stationary phase *Staphylococci* and *Corynebacterium* grown in microaerophilic conditions at pH 5.5 (similar to the human skin^26^), as a representative of volatile production in a skin like environment. We collected 39 strains of publically available skin commensal bacteria (20 *Staphylococci* and 19 *Corynebacterium)* (Fig. 1). Interestingly, we found lactic acid and acetic acid to be the most abundantly produced volatile in both *Staphylococci* and *Corynebacterium* cultures (Fig. 1). Lactic acid is the most well described mosquito attractant emanating from humans ^27^, with nearly all of the highest producers found in the *Staphylococci* genera (Fig. 1). Of the odorants quantified, we chose to test lactic acid and acetic acid, for their high abundance (Fig. 1), and 2-methyl butyrate due to their known impact in mosquito short range behavior^10^, for further mosquito behavior evaluation.

**Figure 1.**
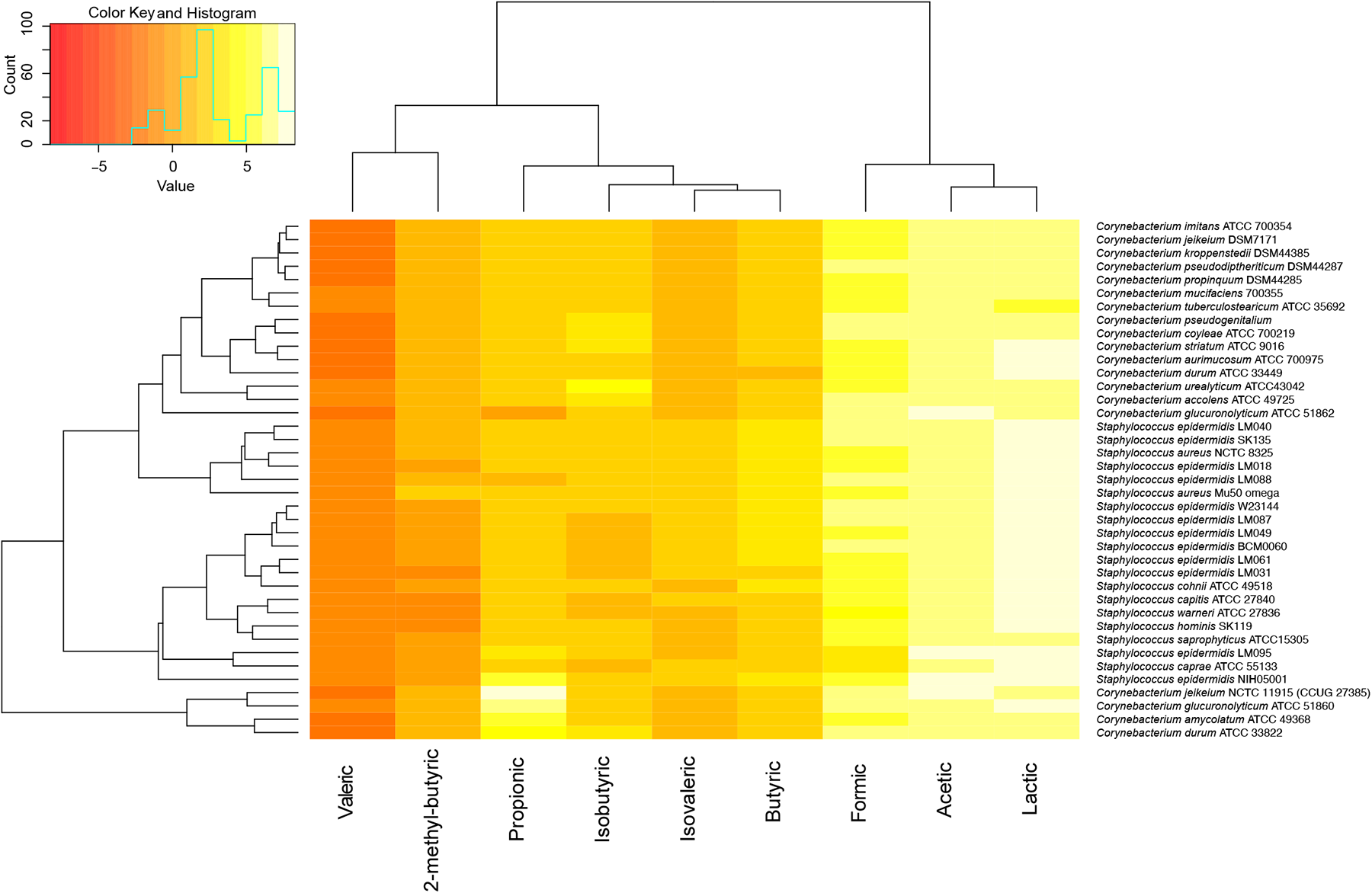
Profiling volatiles produced by human skin bacteria. Heatmap depicting the absolute amounts of specific odorants produced by different strains of *Staphylococcus sp.* and *Corynebacterium sp.* Scale represents log2 values of the concentration in ⲙM.

### Skin odorants synergize to evoke mosquito landing behavior

In order to evaluate the potential of chemical volatiles to reduce mosquito landing behavior, we set up a behavioral arena (Fig. 2A) where female *A. aegypti* had a choice between meshes coated with different odorants placed on opposite sides of the cages (Fig. 2A). Mosquito activity (time spent on each side of the experimental arena) was tracked and recorded using animal tracking at millisecond resolution (Fig. 2B). Carbon dioxide was applied in all experiments. Mosquito landing behavior was first evaluated against the human skin produced odorant and known mosquito attractant L-(+)-lactic acid ^27^(Fig 2C and Suppl. Fig. 1A). Mosquitoes showed stronger attraction to L-(+)-lactic acid at 0.1% (88.9% attraction; Suppl. Fig. 1A) and 0.05% (84.3% attraction; Fig. 2C) than to the water-coated mesh. Mosquitoes still showed attraction at 0.001% (86.2% attraction; Suppl. Fig. 1A) and repellency at the lowest concentration tested (0.0001%, 75.4% repellency; Suppl. Fig. 1A).

**Figure 2.**
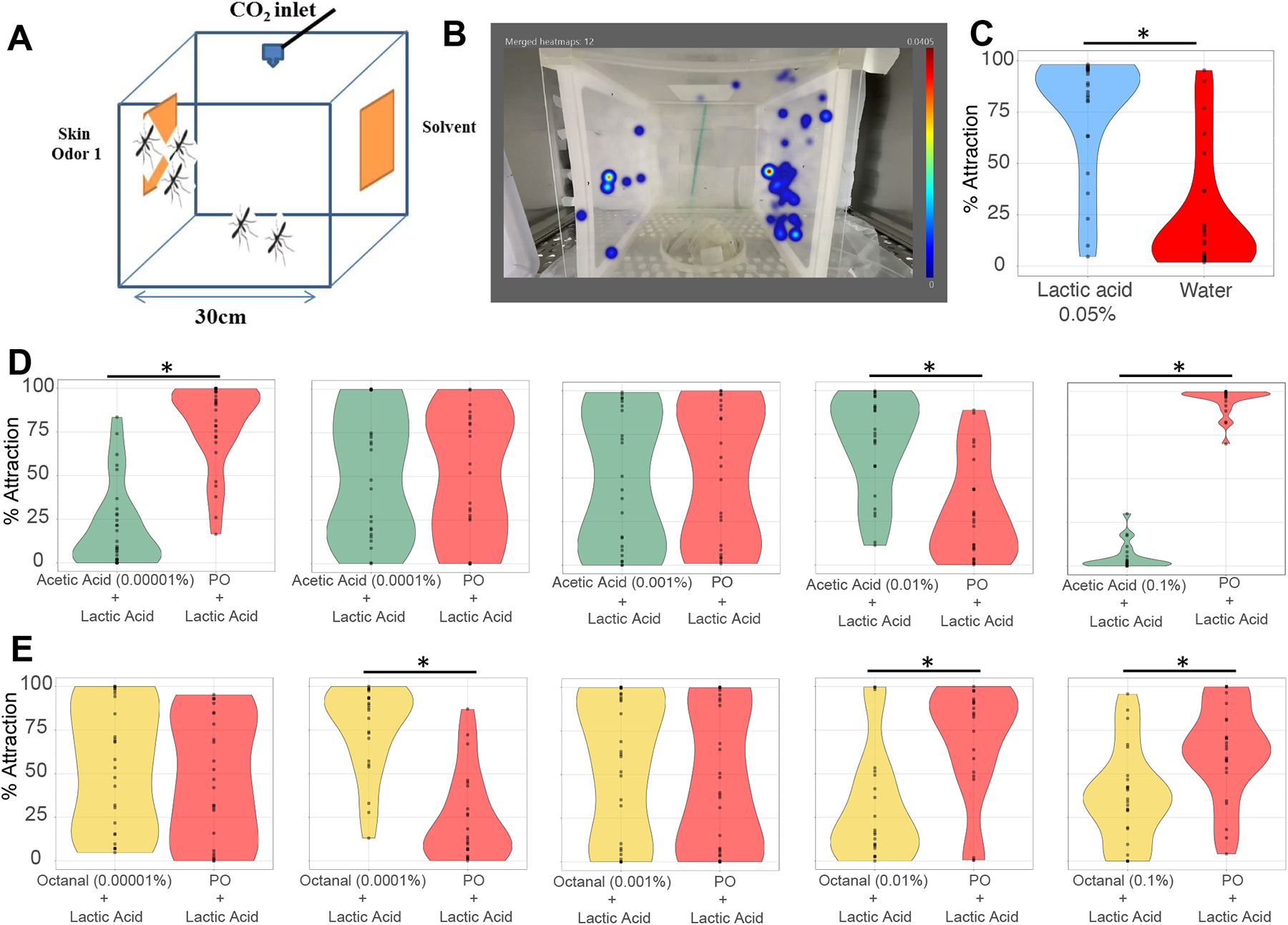
Mosquito 2-choice landing assay. **A.** Schematic representation of a mosquito experimental cage, depicting the odorant-coated meshes on the sides in yellow color, and the carbon dioxide (CO_2_) outlet on the top of the cage in blue color. **B.** Representative picture of a heatmap analysis obtained with the EthoVision software showing the cumulative duration in blue color of mosquitoes on each side of the experimental cage. **C.** Violin plot showing the cumulative duration of the time spent by mosquitoes on the sides of the cages treated with either L-(+)-lactic acid (0.05%) or water. **D-G.** Dose-response assays demonstrating the behavioral responses of mosquitoes to overlays of L-(+)-lactic acid and a skin odorant versus L-(+)-lactic acid and paraffin oil as a solvent. Whereas L-(+)-lactic acid was tested at 0.05% across all experiments, the other skin odorants were assessed at 0.00001%, 0.0001%, 0.001%, 0.01%, and 0.1%. The skin odorants assessed were acetic acid (**D**) and octanal (**E**). Statistically significant differences at p < 0.05 are indicated by an asterisk (*). n = 2 biological replicates, for which the behavior activity of individual mosquitoes was recorded and represented by each dot. Plots represent pooled data of the biological replicates.

As short range y-tube olfactometer experiments indicate that L-(+)-lactic acid along with carbon dioxide gates mosquito attraction and synergize with other skin volatiles ^28–30^, we evaluated mosquito landing behavior against other skin volatiles in the presence of L-(+)-lactic acid at 0.05% and carbon dioxide (Figs. 2D-G and Suppl. Fig. 1B). Acetic acid, a known human skin volatile ^22^, evoked mosquito attraction or repellency in a concentration dependent manner (Fig. 2D). Whereas at the highest and lowest concentrations, acetic acid evoked repellency behavior (97.4% and 89.6% repellency, respectively), this odorant triggers attraction at 0.01% (66.2% attraction; Fig. 2D). Similarly, octanal, another skin volatile ^22,31^, evoked repellency at the two highest concentrations tested (46.5% and 80.6% repellency), but induced landing at 0.0001% concentration (85.1% attraction; Fig. 2E). The evaluation of mosquito landing behavior in the presence of acetic acid or octanal but in the absence of L-(+)-lactic acid resulted in little to null odor induced behavior (Fig. 3). Altogether, these experiments demonstrated that the synergism between skin odorants and L-(+)-lactic acid and carbon dioxide is also applied in a landing behavior context.

**Figure 3.**
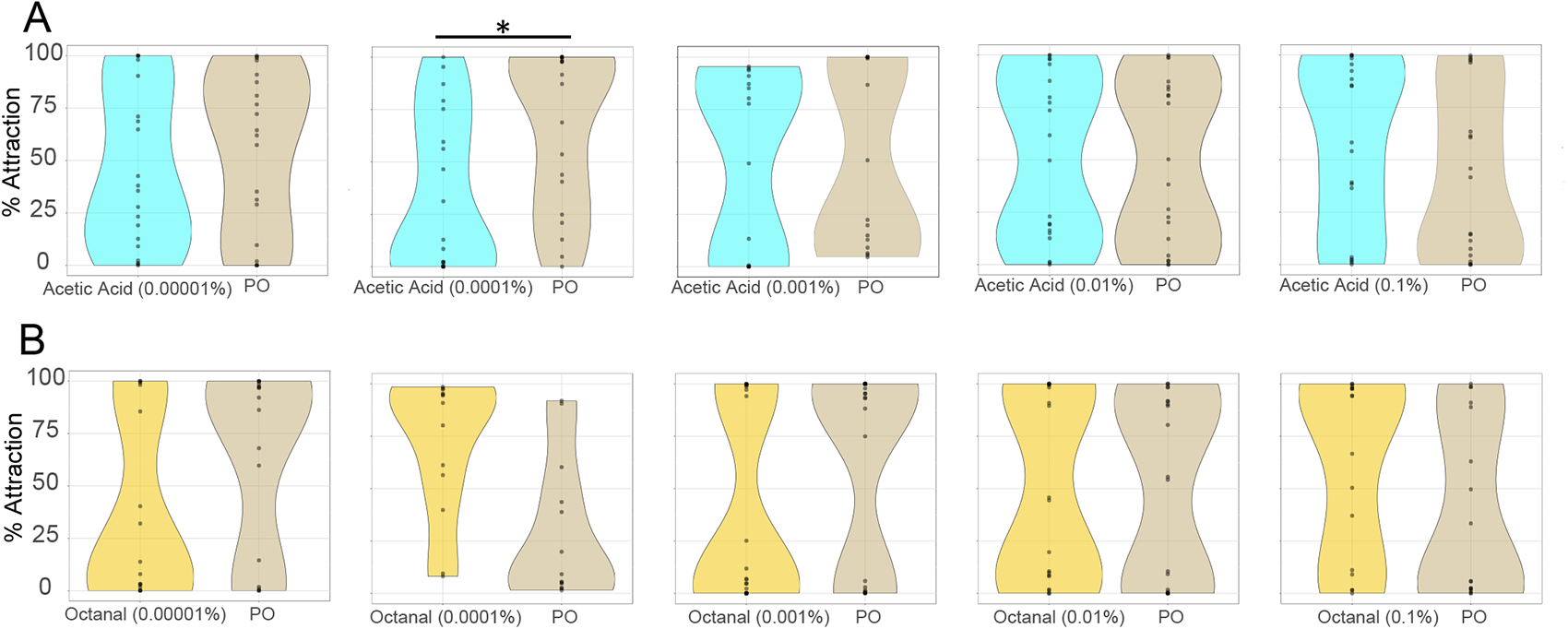
Landing responses against specific skin odorants in the absence of L-(+)-lactic acid. Dose-response assays were performed, as shown in Figure 1, at five different concentrations, using acetic acid (**A**) and octanal (**B**) as testing odorants. Statistically significant differences at p < 0.05 are indicated by an asterisk (*). n = 2 biological replicates, for which the behavior activity of individual mosquitoes was recorded and represented by each dot. Plots represent pooled data of the biological replicates.

### Odorants that reduce mosquito landing behavior

Another odorant isolated from human sweat, 3-methyl butyric acid ^29^ has been shown to induce repellency ^29^ or be inert ^32^ contingent upon the assay used (y-tube olfactometer or traps). This odorant, along with another skin bacteria volatile^33^ structurally similar (2-methyl-butyric acid), was evaluated for landing behavior in the presence of carbon dioxide and L-(+)-lactic acid (Figs 4A and B). In a landing behavior context, 2-methyl butyric acid induced attraction at the highest concentration tested (56.4% attraction; Fig. 4A) but acted as a repellent at all the other concentrations tested (62.0-81.6% repellency range; Fig. 4A). Despite their similar chemical structures, 3-methyl butyric acid evoked repellency at the three highest concentrations (87.1-99.6% repellency; Fig. 4B), but it was inert at the other two concentrations (Fig. 4B).

**Figure 4.**
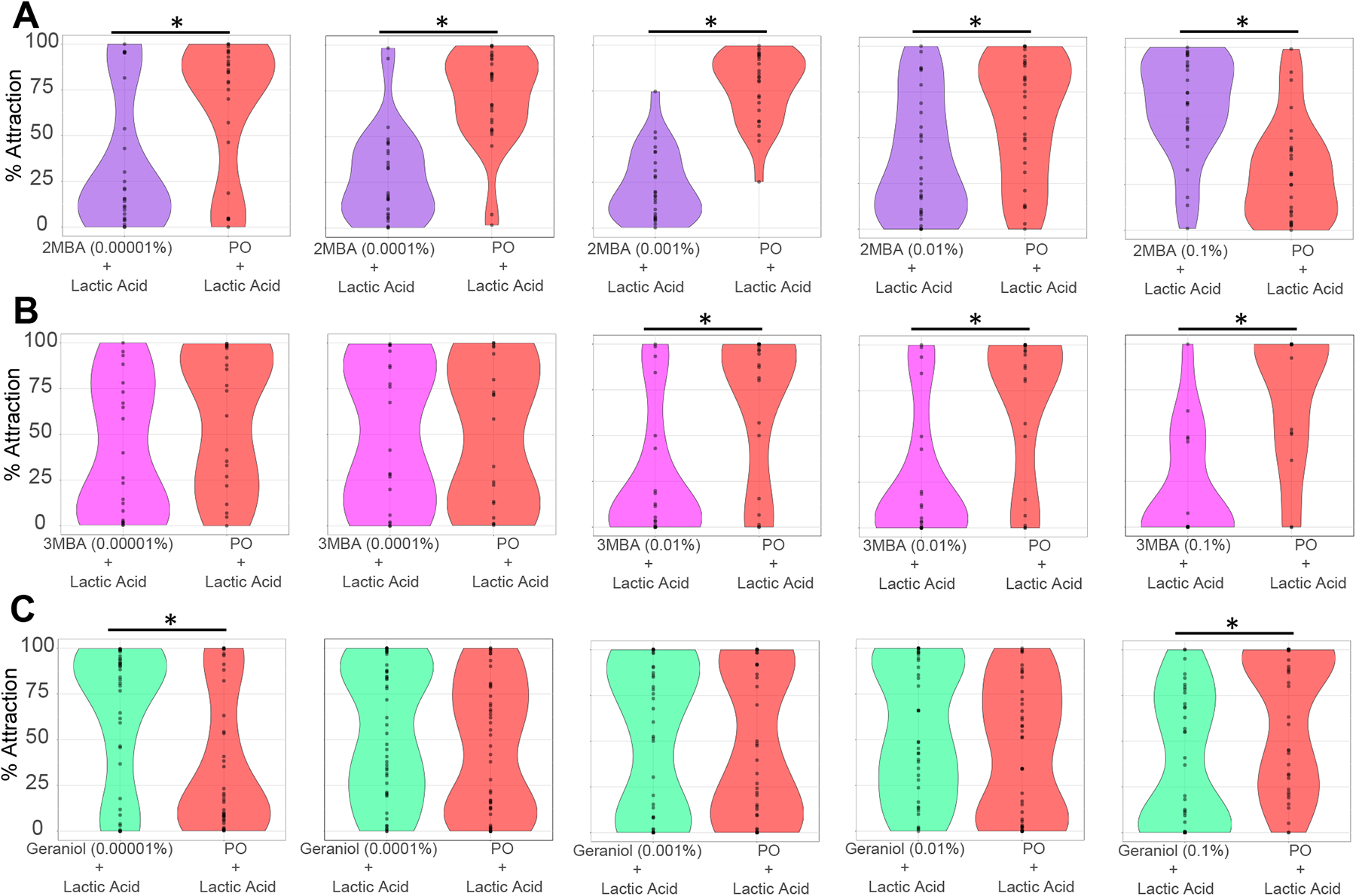
Assessing odorants as repellents for mosquito landing. Dose-response assays were performed as shown in Figure 1, at five different concentrations, using 2-methyl butyric acid (**A**), and 3-methyl butyric acid (**B**), the terpene geraniol (**C**) as the tested odorant overlaid with L-(+)-lactic acid. Statistically significant differences at p < 0.05 are indicated by an asterisk (*). n = 2-3 biological replicates, for which the behavior activity of individual mosquitoes was recorded and represented by each dot. Plots represent pooled data of the biological replicates.

After demonstrating 2-methyl- and 3-methyl butyric acids evoked consistent repellency behavior in the landing context (Figs 4A and B), we evaluated whether other natural odorants could also prevent mosquito landing. As terpenes are commercialized as mosquito repellents ^34^, we assessed if geraniol could also reduce mosquito landing behavior in the presence of carbon dioxide and L-(+)-lactic acid. Geraniol induced landing at the lowest concentration tested (77.7% attraction; Fig. 4C) and repelled mosquitoes from landing on L-(+)-lactic acid-coated mesh at the highest concentration tested (74.9% repellency; Fig. 4C). These experiments indicated that geraniol could also be used as a mosquito repellent.

### Skin volatiles synergize with L-(+)-lactic acid and carbon dioxide in short range attraction behavior

Even though geraniol reduces mosquito landing behavior, other synthetic repellents like DEET and picaridin are effective at preventing mosquito landing ^35^. However, the low volatility (vapor pressure) of these synthetic repellents prevents them from acting effectively at short range ^12^. Terpenes, on the other hand, exhibit higher volatility than synthetic repellents ^35^, which can potentialize the repellency effects provided by the topical application of synthetic repellents. In order to assess the potential of geraniol as short range mosquito repellents, we used a 4-port-olfactometer ^36^(Fig. 5A) that allows mosquitoes to perform most (if not all) host-seeking behavior steps, such as activation, up-wind flight, orientation, and landing (near but not on the odorant source).

**Figure 5.**
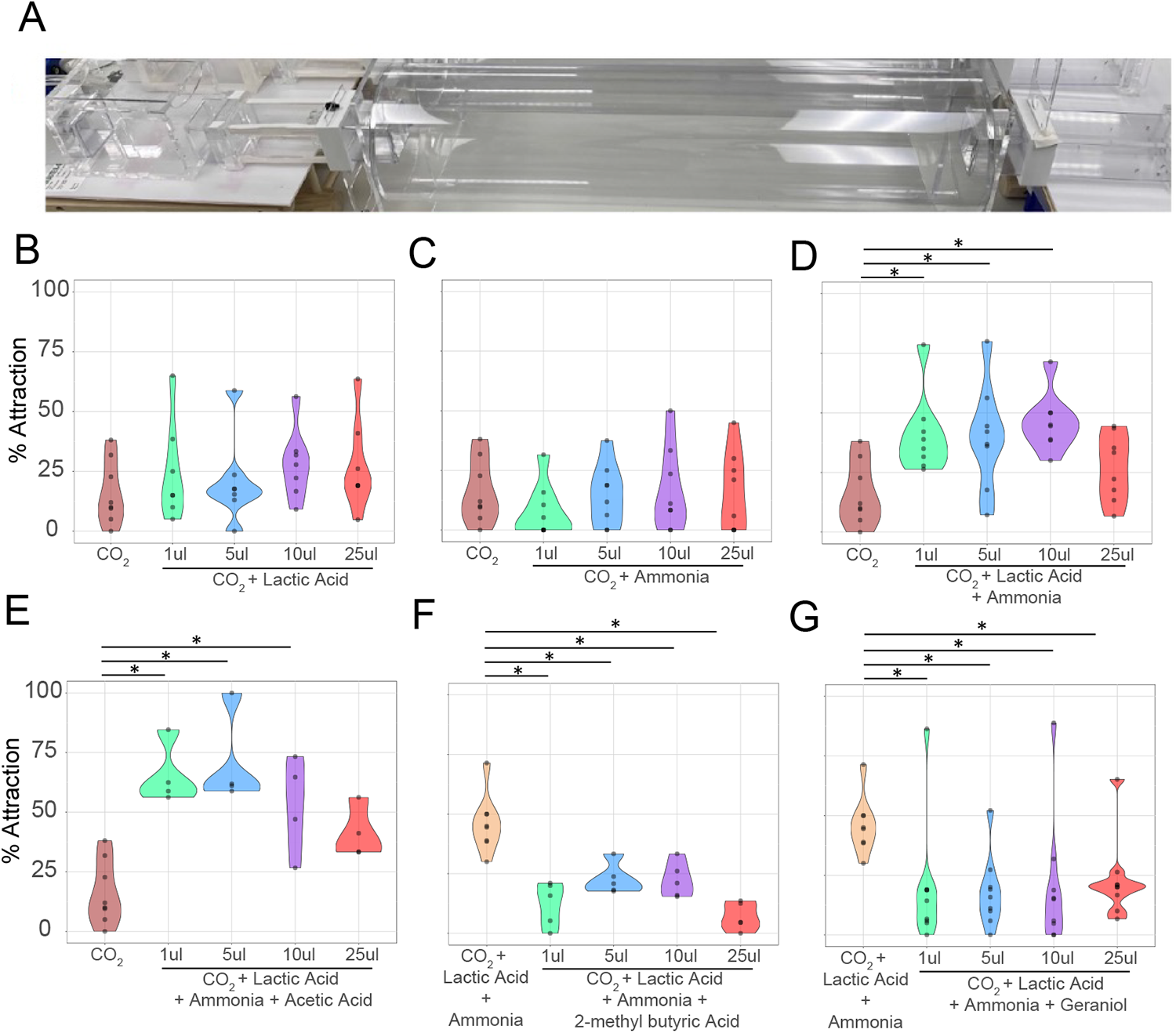
Mosquito short range behavioral assays using a 4-port olfactometer. **A.** Picture depicts a side view of one port of the olfactometer showing from right to left the releasing canister, the flight tube, the trap, and the odorant box. Air flows from left to right. **B-G.** Dose-response behavior assays testing skin odorants and geraniol at 1% concentration and four different doses (1ul, 5ul, 10ul, and 25ul). Attraction to carbon dioxide (CO_2_) alone and CO_2_, L-(+)-lactic acid, and ammonia were used as standards for attraction (**B-E**) and repellency (**F-G**) assays, respectively. Such standards’ data are replicated in each graph and represent a single experiment. **B.** Mosquito attraction to either CO_2_ alone or CO_2_ along with 4 doses of L-(+)-lactic acid along with CO_2_. **C.** Mosquito attraction to either CO_2_ alone or CO_2_ along with 4 doses of ammonia acid along with CO_2_. **D.** Behavioral responses to CO_2_, L-(+)-lactic acid, and different doses of ammonia for mosquito attraction. L-(+)-lactic acid was tested at 5ul dose. **E**. Mosquito attraction to combinations of CO_2_, L-(+)-lactic acid (5ul), ammonia (10ul), and different doses of acetic acid. **F-G.** Behavioral responses of mosquitoes to combinations of CO_2_, L-(+)-lactic acid (5ul), and ammonia (10ul), and different doses of 2-methyl butyric acid (**F**), and geraniol (**G**). Statistically significant differences at p-adjusted < 0.05 are indicated by an asterisk (*). n = 4-9 biological replicates. Each dot represents the percentage of mosquitoes caught in the olfactometer traps for each biological replicate.

In order to demonstrate that the 4-port olfactometer can be used to assess short range mosquito behavior and establish positive controls for attraction and repellency, we assessed different combinations of human skin odorants (Fig. 5A-E). Initially, we tested mosquito attraction to four different doses of either L-(+)-lactic acid (Fig. 5B) or ammonia (Fig. 5C) at 1% concentration in the presence of carbon dioxide. The presence of such odorants alone did not evoke a statistically significant improvement in mosquito attraction compared to carbon dioxide alone (Figs. 5B and C). In contrast, combinations of L-(+)-lactic acid (5 µl) and three different doses of ammonia increased mosquito attraction when compared to carbon dioxide alone (40.3-64.9% attraction improvement; Fig. 5D). Addition of acetic acid at three different doses to a combination of L-(+)-lactic acid (5 µl) and ammonia (10 µl) synergistically improved mosquito attraction (60.6-77.1% attraction improvement; Fig. 5E). These findings corroborate previous studies using y-tube olfactometers ^30^, and validated a blend of L-(+)-lactic acid (5 µl), ammonia (10 µl), and carbon dioxide as an human-derived attractive cue for the following mosquito repellency assays.

### Geraniol reduces mosquito attraction at short range

As 2-methyl butyric acid was shown to consistently prevent mosquito landing at multiple concentrations (Fig. 2F), we first assessed whether 2-methyl butyric acid could also evoke mosquito repellency in the 4-port olfactometer. This odorant significantly reduced mosquito attraction to carbon dioxide, L-(+)-lactic acid, and ammonia at all doses (Fig. 5F). As geraniol also reduced mosquito landing behavior (Fig. 4C), we assessed the potential of this terpene to reduce mosquito attraction at short range. Geraniol showed statistically significant reduction in mosquito attraction at all doses when compared to carbon dioxide, L-(+)-lactic acid, and ammonia (69.2-77.9% repellency range, Fig. 5G).

## Discussion

The human skin is covered with different sweat glands that are localized in different areas of the body^23^. Whereas the eccrine glands are distributed all over the body, apocrine glands are localized in the moist regions of the body (groin and axilla), and sebaceous glands are more localized to the face and torso (sebaceous areas; ^23^). Different bacteria species of the human resident skin microbiome are associated with such glands, which release different types of biomolecules used by the bacteria as nutrients ^10,26^. Upon the metabolism of these nutrients, the small molecules released are highly attractive to mosquitoes ^37^.

The behavioral effects of the human kairomones carbon dioxide, L-(+)-lactic acid, and ammonia together gating mosquito short-distance attraction and trap catching for both *Anopheles gambiae* ^38,39^ and *A. aegypti* ^7,30^ mosquitoes have been well established. In the absence of L-(+)-lactic acid and/or ammonia, mosquito attraction to carbon dioxide is not induced by the other carboxylic acids secreted by the human skin ^7,27,38^. In this study, we demonstrated that a similar principle also governs landing behavior, as landing behavior to specific human skin odorants such as acetic acid and octanal was abrogated in the absence of L-(+)-lactic acid. Our findings also pointed out that specific skin odorants, such as 3-methyl butyric acid, can also repel mosquito landing behavior, as it has been demonstrated for short distance attraction ^29,38^. We have also assessed the effects of the terpene geraniol in the context of short range attraction to a blend of human kairomones, which was capable of reducing mosquito attraction.

Altogether, these findings point to multiple targets of the skin microbiome that can be genetically manipulated to reduce the synthesis of important odorants that govern mosquito landing behavior. The biosynthetic pathway that synthesizes L-(+)-lactic acid stands as a main target, as this odorant is produced at very high levels by the human skin bacteria, and we and others ^7,30^ have shown that this odorant gates (along with carbon dioxide) the mosquito short range attraction and landing behaviors evoked by other skin odorants. Knocking down the synthesis of ammonia seems also to be a good strategy as this odorant is even more important than L-(+)-lactic acid to gate behavioral responses of the mosquito *Anopheles coluzzii* ^38,39^. Another interesting target is a gene associated with the acetic acid-producing pathway, as this odorant is also produced at high levels by skin microbes and synergizes the mosquito behavioral responses triggered by L-(+)-lactic acid and ammonia. Alternatively, using genetic tools to induce the synthesis of repulsive odorants might potentially reduce mosquito bites. Making the human scent unattractive has the potential to divert anthropophilic mosquitoes to feed upon other animals, reducing pathogen transmission and disease burden.

## Materials and Methods

### Culturing skin commensal bacteria

All bacterial strains were stored at −80℃ in 25% glycerol until experimentation. To grow strains for GC/MS quantification, strains were first plated on BHI + 1% Tween agar plates (1.5% agar) and grown for 1 (*Staphylococci*) or 2 days (*Corynebacterium*) aerobically at 37℃, at which point approximately 1µL of cell material was transferred into 10mL prewarmed BHI + 1% Tween at pH 5.5 broth in a 15mL conical tube. The conical tube was screwed tight and cultures were incubated at 37℃ for 1 (*Staphylococci*) or 2 (*Corynebacterium*) days to reach stationary phase growth.

### GC/MS analysis of skin commensal cultures

For analysis of commensal supernate, one ml of stationary phase bacterial culture was centrifuged at 13,000 x g for 10 min at room temperature. 400 µl of extraction solution (20 µL 10 mM n-crotonic acid in water as internal standard, 100 µL 6 N HCl, 280 µL ddH_2_O), 100 µl of cell-free supernatant, and 500 µL diethyl ether were added together in beads tube. In parallel, standards were created to facilitate quantification by adding 100 µl of SCFAs mix solution (ranging from 5,000 µM to 0.5 µM, series of half dilution) into 400 µl of extraction solution (20 µL 10 mM n-crotonic acid in water as internal standard, 100 µL 6 N HCl, 280 µL ddH_2_O) and 500 µL diethyl ether in beads tube.

Using a QIAGEN Tissue Lyser II, samples were mixed at 25/s for 10 min. The resulting homogenates were subjected to centrifugation at 18000 x g for 10 min, organic layer, and transferred to a new glass vial (29391-U, Supelco) for derivatization. This was achieved by first taking 100 µl of diethyl ether extract and mixing with 10 µL MTBSTFA and incubated at room temperature for 2 h. 1 µL of the derivatized samples were analyzed using a 7890B GC System (Agilent Technologies), and 5973 Network Mass Selective Detector (Agilent Technologies). Derivatized samples were analyzed using the following chromatography conditions for GC-MS: Column: HP-5MS, 30 m, 0.25 mm, 0.25 µm; Injection Mode: splitless; Temperature Program: 40 °C for 0.1 min; 40-70 °C at 5 °C/min, hold at 70 °C for 3.5 min; 70-160 °C at 20 °C/min; 160-325 °C at 35 °C/min, equilibration for 3 min. 1 µL of each sample was injected and analyte concentrations were quantified by comparing their peak area standards created using pure representatives.

### Synthetic chemical volatiles and odorant dilutions

Synthetic odorants were purchased from Sigma-Aldrich unless otherwise specified at the highest purity. Odorants were diluted in either molecular grade water or paraffin oil (PO) to 1% v/v before use.

### Mosquito maintenance and starvation

*Aedes aegypti* Liverpool strain mosquitoes were raised and maintained according to ^40^. Seven to twenty-one days old nulliparous females were sorted into groups of 25 specimens, transferred to the releasing canister of the olfactometers (described below), and starved for 5-8 hours without water at 28℃ and 70% relative humidity (RH).

### Mosquito behavioral assay - 2-choice landing assay

Mosquito landing assays were performed in Bugdorms (30×30×30 mosquito cages) inside a mosquito incubator (Caron, Marietta OH). For filming, one of the sides of the cage was replaced by a transparent plastic pane secured with white Duck tape. Odorants or solvents (water or PO) were applied (600 μl) onto white polyester nets (10×10cm; Bioquip CAT#7250A) laying on a glass Petri dish and hanging onto the opposite lateral side of the bug dorm using push pins. For overlay experiments, L-(+)-lactic acid-coated mesh along with another mesh coated with the tested odorant were hung on the experimental cage with the L-(+)-lactic acid mesh in contact with the cage. On the opposite side, an L-(+)-lactic acid-coated mesh was hung along with a solvent-coated mesh. The positions of the control mesh and the tested odorant mesh were switched amongst trial replicates. Pure carbon dioxide was delivered using a fly pad placed face down onto the experimental cages.

On the day before the experiments were performed, 16 mated nulliparous mosquitoes were transferred to individual bug dorms and starved overnight with deionized water. The behavior trials were carried out on the next day between 1-5 pm, and videos were recorded for 5 minutes after the first minute upon switching the carbon dioxide regulator on.

### Mosquito behavioral assay - high-throughput (HT) olfactometer

Short range mosquito behavioral assays were performed with the 4-port high-throughput olfactometer ^36^. Room temperature and humidity were maintained at 27.5℃ and 60% relative humidity using space heaters and humidifiers. Purified air was pumped into the system at 24,367 mL/min rate, whereas pure CO_2_ was flown at 254 mL/min (final concentration per lane ∼ 1,500-2000 ppm). Starved mosquitoes were exposed to air only for 10 min, when odorants and/or bacterial cultures were placed in the odor chamber onto 47mm plastic Petri dishes (Fisherbrand), and CO_2_ gauge was switched on. The gates of the releasing canisters were open, and the behavioral assays were carried out for 20 min. Then, both the releasing canister and the trap gates were closed, and the number of mosquitoes in the releasing canisters, flight tubes, and traps were scored. The tested odorants and cultures were switched amongst the 4-port olfactometer across trial replicates. Dose-response assays were carried out to determine the doses of chemicals and/or bacterial cultures that evoked the strongest behavioral responses. Doses of 1 μl, 5 μl, 10 μl, 25 μl were tested.

### Video recording of behavioral activity

For the 2-choice assay, videos of mosquito activity were recorded with an iPhone X at 30 fps. Videos were then analyzed with the EthoVision XT software (Noldus) at millisecond resolution and using individual mosquito tracking. Only experiments whereby at least 40% of the mosquitoes were active and tracked were analyzed.

### Behavior apparatus cleaning

All equipment used in behavior assays was soaked overnight (small parts) or washed thoroughly (flight tubes) with scent-free laundry detergent (Seventh Generation, free & clear) and rinsed with tap water thoroughly.

### Statistical analyses

Graphs and statistical analyses were performed with the R software. For both 2-choice landing and 4-port olfactometer experiments, time spent on each side of the experimental cages and the number of mosquitoes caught by the traps were transformed into percentages so as to normalize for mosquito participation variability across experimental replicates. Shapiro-Wilk normality test was used to assess whether or not the data fit a normal distribution. For pairwise comparisons, either the Welsh t-test or Wilcoxon rank sum test were used. For multiple comparisons, either ANOVA or Kruskal-Wallis’s rank sum test were applied. These tests were followed by post-hoc analyses using Tukey multiple comparisons of means and Wilcoxon rank sum test, respectively. p-values were adjusted (p-adjusted) for multiple comparisons using the Benjamini-Hochberg procedure. All raw and analyzed data can be found in the **Supplementary Table S1**.

## Acknowledgments

We thank Judy Ishikawa and Ava Stevenson for helping with mosquito husbandry and Rhodri Edwards for critically reading the manuscript. This work was supported by funding from a DARPA contract (HR00112020030) awarded to O.S.A and M.F. and NIH grants (RO1AI148300, RO1AI175152, RO1AI151004) awarded to O.S.A. The views, opinions, and/or findings expressed are those of the authors and should not be interpreted as representing the official views or policies of the U.S. government.

## Author Contributions

O.S.A. and M.A.F. conceptualized the project. IC-A and JAM designed the experiments. IC-A and OJ carried out behavior assays. KA performed GC-MS experiments. IC-A, RR, and JAM analyzed and compiled the data. All authors contributed to the writing and approved the final manuscript.

## Ethical conduct of research

All animals were handled in accordance with the Guide for the Care and Use of Laboratory Animals as recommended by the National Institutes of Health and approved by the UCSD Institutional Animal Care and Use Committee (IACUC, Animal Use Protocol #S17187) and UCSD Biological Use Authorization (BUA #R2401).

## Disclosures

O.S.A is a founder of Agragene, Inc. and Synvect, Inc. with equity interest. The terms of this arrangement have been reviewed and approved by the University of California, San Diego, in accordance with its conflict of interest policies. All other authors declare no competing interests.

## Data availability

All data generated or analyzed during this study are included in this published article and its supplementary information files.

## Supplemental Figure Legend

**Supplemental Figure S1.**
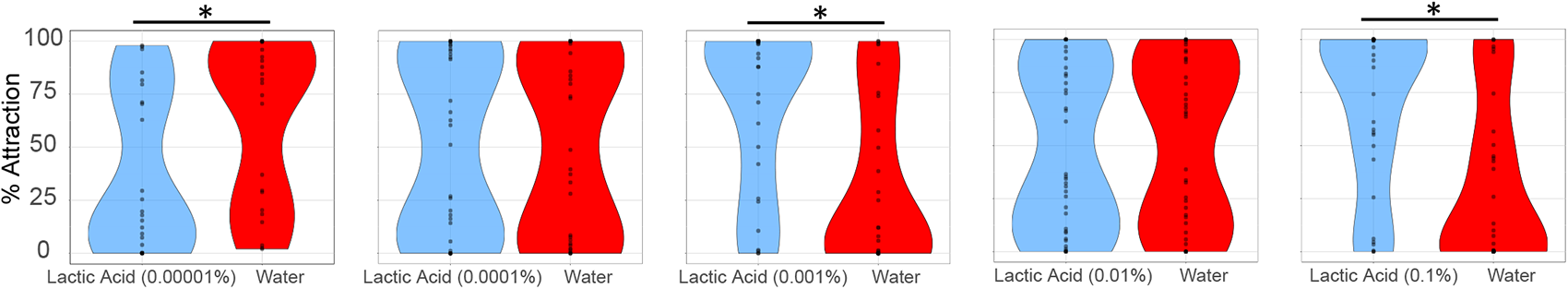
Two-choice landing assays with skin odorants. Dose-response assays were performed, as shown in Figure 1, between three and five different concentrations, using the L-(+)-lactic acid as testing odorant as the tested odorant. Statistically significant differences at p < 0.05 are indicated by an asterisk (*). n = 3 biological replicates, for which the behavior activity of individual mosquitoes was recorded and represented by each dot. Plots represent pooled data of the biological replicates.

## Notes

### Summary of Updates

This version has been corrected to focus the text more on the hypothesis tested and to fix a misplaced statistical bar in Fig 2

